# Hermes: an ensemble machine learning architecture for protein secondary structure prediction

**DOI:** 10.1101/640656

**Authors:** Larry Bliss, Ben Pascoe, Samuel K Sheppard

**Affiliations:** The Milner Centre for Evolution, Dept. of Biology and Biochemistry, University of Bath, UK

## Abstract

**Motivation:** Protein structure predictions, that combine theoretical chemistry and bioinformatics, are an increasingly important technique in biotechnology and biomedical research, for example in the design of novel enzymes and drugs. Here, we present a new ensemble bi-layered machine learning architecture, that directly builds on ten existing pipelines providing rapid, high accuracy, 3-State secondary structure prediction of proteins.

**Results:** After training on 1348 solved protein structures, we evaluated the model with four independent datasets: JPRED4 - compiled by the authors of the successful predictor with the same name, and CASP11, CASP12 & CASP13 - assembled by the Critical Assessment of protein Structure Prediction consortium who run biannual experiments focused on objective testing of predictors. These rigorous, pre-established protocols included 7-fold cross-validation and blind testing. This led to a mean Hermes accuracy of 95.5%, significantly (p<0.05) better than the ten previously published models analysed in this paper. Furthermore, Hermes yielded a reduction in standard deviation, lower boundary outliers, and reduced dependency on solved structures of homologous proteins, as measured by NEFF score. This architecture provides advantages over other pipelines, while remaining accessible to users at any level of bioinformatics experience.

**Availability and Implementation:** The source code for Hermes is freely available at: https://github.com/HermesPrediction/Hermes. This page also includes the cross-validation with corresponding models, and all training/testing data presented in this study with predictions and accuracy.

## Introduction

Over 65 years of protein structure prediction research has been accompanied by progress in X-ray crystallography (X-RC), nuclear-magnetic resonance, cryo-electron microscopy, and volumetric electron microscopy. Protein structure prediction is now an important component of biotechnology and biomedical research, for example in the design of novel enzymes and drugs. Experimental studies have yielded around 132,000 entries in the Protein Data Bank (PDB), with about 10,000 additions every year. However, for an estimated cost of $100, 000 per protein (including cloning, expression, purification, and structure determination) only around 4.6% of analyses yield solved structures and PDB entries [1]. Therefore, there is a great incentive to develop rapid, accurate and inexpensive tools for protein prediction to improve drug design, enzyme active site detection, and allow greater mechanistic understanding of the genetic basis of phenotype variation.

Early protein structure predictions estimated the folding propensities of amino acids based on data-mining of the just 29 protein structures that had been experimentally determined [2]. This approach was surpassed by successive developments using GOR algorithms that utilised a sliding window and Bayesian conditional probabilities [3–7]. Prediction accuracy varied, at around 70%, until the introduction of rapid multi-sequence alignments (MSAs) to capture evolutionary information, in concert with machine learning techniques. Today, two major approaches have emerged as the most successful. The first, *ab initio* approach, predicts structure solely from amino acid sequence. The second, template-based prediction, fragments the protein input and searches the PDB database for solved proteins with similar amino acid sequence (Using a BLAST identity threshold) and records the structures as solutions for the query sequence [8,9]. Any regions lacking a homologous match are then predicted using an *ab initio* method. Because template-based methods use large amounts of previously defined information they generally achieve around 7% greater accuracy for any given protein [10,11]. As with gene prediction, the distinction between the methods has blurred [12]. Template-based methods rely on the archive of homologous protein sequences, and every year this improves, with the current probability of any query sequence sharing insufficient homology with existing structures being <5% [13]. Challenges remain in optimising protein prediction approaches, especially at boundary regions between secondary structure elements, due to inconsistent assignment of states coupled with the limited conservation of element length. Moreover, sequence homology does not perfectly predict structural homology and structural homologs can share as little as 20% identity [14]. This indeterminacy, sometimes referred to as a ‘Twilight zone’, and occurs at ≈20-35% sequence identity and, while its existence is widely accepted, the causality is debated [15].

Here we introduce Hermes, an ensemble bi-layered protein secondary structure predictor. This machine learning tool automatically switches between *ab initio* and template-based methods and capitalizes on advances in high performance computing to allow for increased complexity of methods, even with small training sets. The first layer integrates ten well established methodologically divergent prediction pipelines. The second layer incorporates this information for meta-learning, via six original classification strategies that experience folding with the two top performing initial layer predictors, a process in which the results of the previous layer are forwarded to the next. Ensemble methods, such as Hermes, have risen in popularity, along with generally deeper architectures, proving efficacious in recent machine learning competitions [16–18]. Hermes utilises a meta-learning approach called stacking to improve strong learners and explore the entirely new hierarchical secondary structure space [19]. Together, stacking and folding improve accuracy over current methods with a validated prediction accuracy of 95.5%.

## Methods

### Model architecture

Hermes is composed of two layers, named for the types of sequences they take as input (Fig. 1). The initial, ‘Protein’ layer, comprises ten secondary structure predictors. Of these, two were developed in the 1990’s: GORIV, built axiomatically with information theory; and PHD, an early neural network [6,20]. As such, their accuracies are significantly lower than the remaining eight (below), however, they provide informed variability to the Protein layer. Of these other eight: NetSurfP, JPRED4, YASPIN, Spider3, and RaptorX-Property (DeepCNF), are *ab initio*, while pS2, SSpro, and Porter4 are primarily template-based methods [21–28]. Moreover, the predominant protein property that each predictor exploits varies: Spider3 utilises key polypeptide bond angles, RaptorX-Property opts for residue contacts, and pS2, SSpro, & Porter4 exploit homology. The wide machine learning framework of Hermes is supplied by the Protein layer, formed from these ten predictors. All Protein layer predictions are obtained via a requests script, that automatically posts the query protein sequence to the respective web servers, e.g. http://RaptorX-Property.uchicago.edu/StructurePrediction/predict/, and then scrapes the HTML to retrieve the prediction. In the event that any Protein layer prediction severs are not operational, Hermes will automatically drop those and continue to provide predictions with those remaining, albeit less accurately. This is achieved by using the same architecture, trained with the same dataset but with the affected Protein layer predictor removed. Folding continues, with the two most accurate predictors available. However, if Porter4 is one of those affected, the user is notified to try again at a later time. Although, this mechanism was never required during the data collection for this work.

**Fig. 1.**
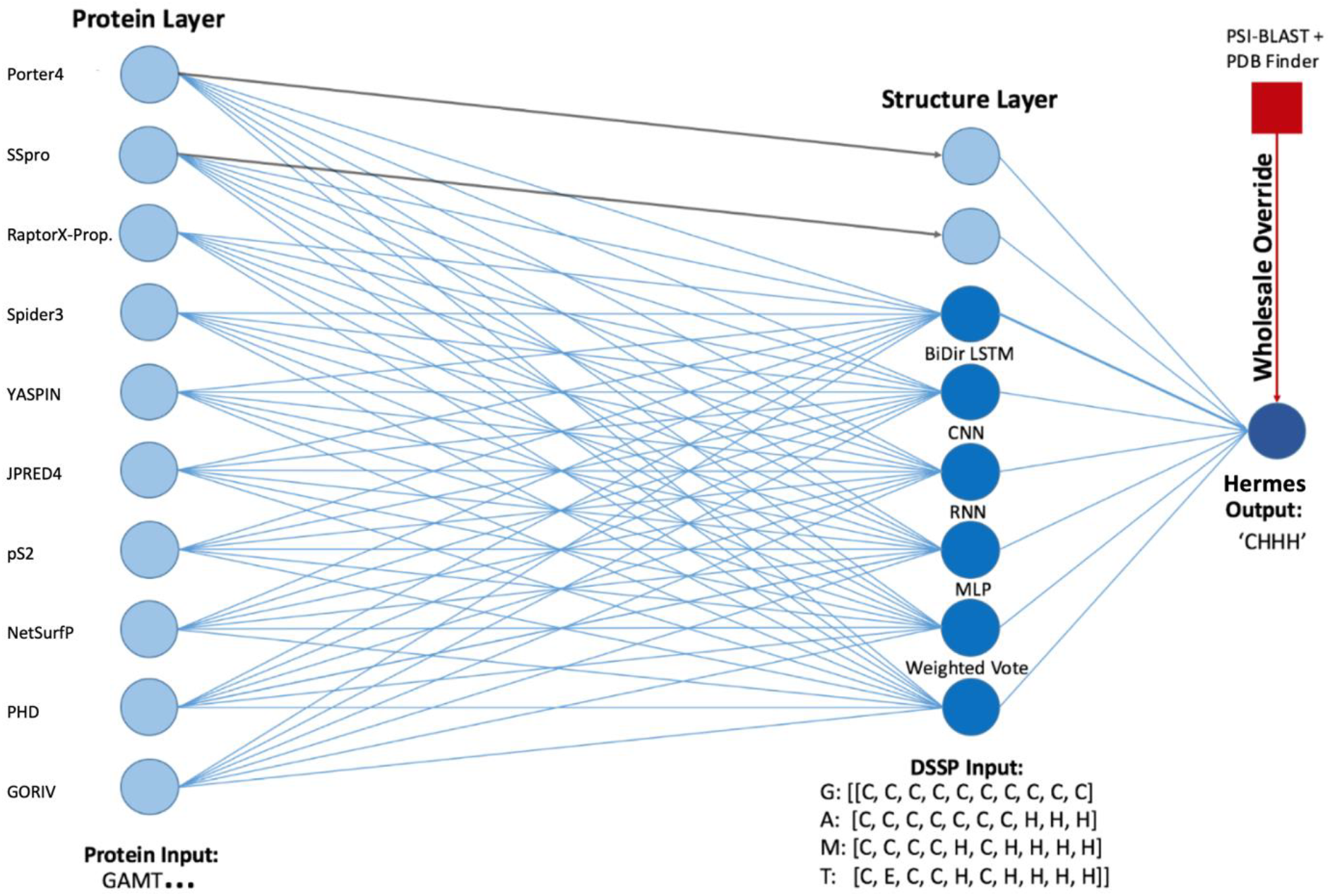
A Schematic of Hermes Architecture. The Hermes design employs two layers: The Protein Layer of 10 predictors, on the left, and the Structure layer of 8 predictors, on the right, into a final aggregator-network to return the final secondary structure prediction. Predictors are identified with circles, the output of which is fed into the next, as shown by blue lines. The model utilises folding, whereby a predictors’ output is both supplied as input to predictors of the next layer and directly carried forward to the next layer. This is performed on the two highest-performing classifiers, SSpro and Porter4, both originate at the Protein layer. Each is displayed with arrows. Sample inputs for each layer are represented under the schematic.

The Structure layer of Hermes comprises eight separate classifiers. Six originate at this layer and individually take all ten Protein layer secondary structure predictions as features to return a refined prediction by meta-learning. These include: a simple weighted vote; a multilayer perceptron (MLP) [29]; a recurrent neural network (RNN) [30]; a stacked gated recurrent unit (GRU) [31]; a stacked bidirectional long short-term memory (BiLSTM) unit [32]; a convolutional neural network (CNN) [33]. Additionally, the outputs of the two highest performing Protein layer predictors, SSpro and Porter4, undergo folding into this second layer. This equates to passing these two predictions twice, once as an input to the other six Structure layer classifiers and then again as an output of the Structure layer. This generates a total of eight high-fidelity Structure layer predictions that are passed through a final aggregator-network, specifically a BiLSTM unit, which gives the final Hermes solution [34]. The output of every element within Hermes, for a single protein, outlines how each plays a role in progressively improving and error correcting input (Fig. 2). This provides the deep learning capabilities of Hermes and the computational load is flattened, via the division of layers, which facilitates local use on standard desktops in the order of minutes.

**Fig. 2.**
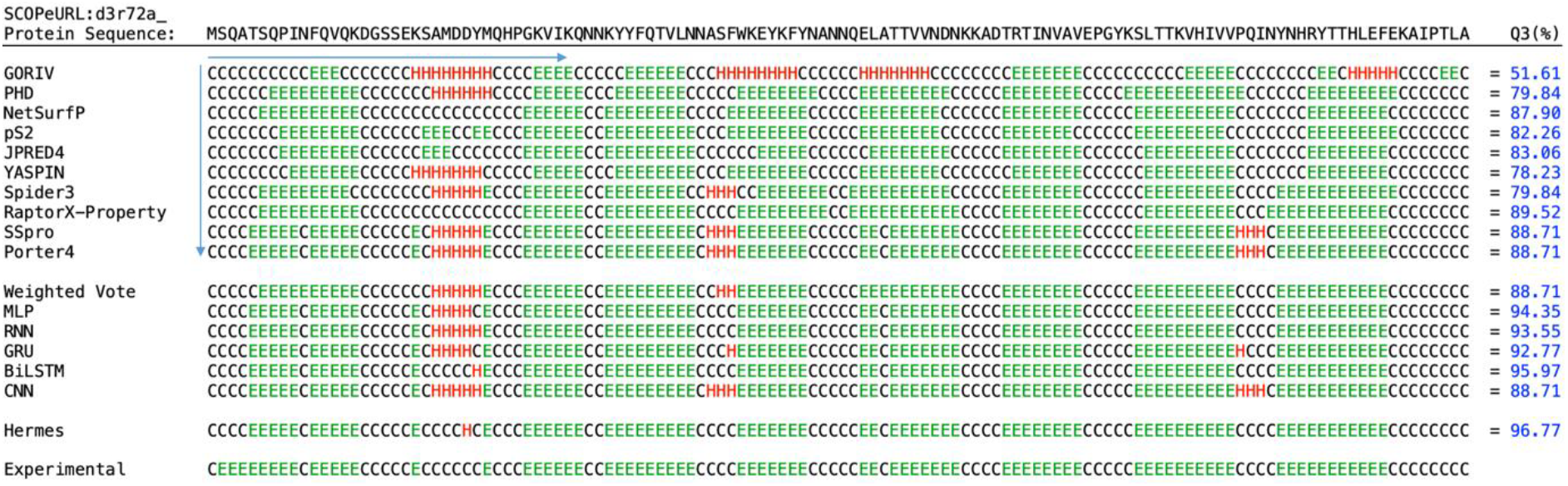
A Sample Output of Every Predictor Used Within Hermes. The output, i.e. secondary structure, of all classifiers within Hermes aligned to a protein, with Q3 values for each shown. DSSP states are identified as: Disordered coil as a black C, alpha-Helices as a red H, and extended beta-Strands as a green E. Vertical and horizontal blue arrows originating from the first prediction by GORIV indicate the information taken in by the CNN, MLP & weighted vote and RNN, GRU & BiLSTM, respectively. Columns identify the prediction of each amino acid by each predictor/classifier and rows show the complete prediction sequence. These predictions are divided into four groups; the Protein layer, the Structure layer, Hermes and the final row is the experimentally determined structure. Predictions are improved as they pass through these groups.

### Training/testing protocol and datasets

For comparability and minimisation of bias, the publicly available JPRED4: JNet v.2.3.1 training and blind-test datasets were used [35]. The former is defined by 1348 protein sequences and the latter by 149, with both being unmodified from the JPRED4 protocol. The training and test sets are derived, and subsequently filtered, from the SCOPe/ASTRAL database and of different structural super families, but all of <2.5Å resolution and pairwise redundant. Only sequences between 30 and 400 residues were used. The training-testing split is stratified, with respect to the three secondary structure states; Alpha-Helix, Beta-Sheet, and disordered coil, within 1% equal composition of each. Further, three more test-only datasets were run, named CASP11, 12 and 13; derived from experiments organised by the Critical Assessment of Techniques for Protein Structure Prediction (CASP) [36,37]. Teams of modellers compete to predict structures, with the targets deliberately selected for difficulty to predict. The list of targets within the “Regular” class are given, with some being cancelled, some failing to provide validation structures, and a further subset providing PDB codes that are not linked to the representative sequence. Hence, for CASP11, 12 and 13 there were 77, 39 and 26 proteins respectively. All four test sets were then redundancy reduced with respect to the training data of all ten protein layer predictors, where such data was used, with a 30% sequence identity cut off [38]. This resulted in a total test set of n=118 proteins: JPRED4 blind test n=48, CASP11 n=40, CASP12 n=20, CASP13 n=10. We ran an initial 7-fold cross validation on the training dataset, followed by training on all 1348 sequences of the training dataset, for a final blind run against the JPRED4 test dataset and the three most recent CASP datasets 11, 12 and 13.

### Model output

The accuracy improvement of the protein sequence as the 11^th^ feature was insignificant, and so not implemented. Hence, the classifiers of the Structure layer, separately, receive a 10-element array. Whereby each element is a single amino acid prediction from a single Protein layer predictor, and the array is the output of the entire Protein layer for that one amino acid. Further, an element is one of the three DSSP states; ‘H’, ‘E’, or ‘C’, e.g. [H, H, H, C, H, C, E, H, C, H]. Each Structure layer predictor then returns a probability matrix of the three labels, whose sum is equal to 1 e.g. [0.2353, 0.7100, 0.0547], due to a final softmax activation function [39].

The information that the Structure layer classifiers are assessing can be divided into two classes. Frist, the RNN, GRU and BiLSTM are analysing each prediction along its length. Second, the MLP, weighted vote and CNN are solving relationships between predictions, given in each 10-element array. This dichotomy of methods and the variety of neural network models within the Structure layer facilitate deep spatial learning. This can, however, result in overfitting, with high accuracy on training data but reduced generalisability [40]. To account for this, each neural network of the Structure layer was trained on 250 epochs, using either nAdam or Root Mean Squared Propagation (RMSprop) optimisers. This measure was integrated in the model with two (or more) of the following protocols: early stopping; dropout; L1/L2 activity regularisation [41–43]. Grid-searches, measured by loss, were used concurrently for optimisation of each model in hyperparameter space [44] for: batch size; hidden layer sizes; initializers; optimisers; and early stopping conditions. Structure layer neural networks are implementations of Googles’ TensorFlow API [45], with Hermes being written entirely in Python.

Finally, if Hermes receives perfect predictions from the Protein layer, which can occur if the query protein has already been solved and exists in the PDB (or an almost identical homolog), the final output maybe no better or even worse. Hermes implements a ‘Wholesale Override’ feature to rectify this. Specifically, a PSI-BLAST search of the entire query sequence is run against the PDB database at E value = 0.001 and identity > 95.5% (the mean accuracy of Hermes). The hit that shares greatest identity is then substituted for the output.

### Evaluation metrics

Consistent with current protein prediction models, Hermes describes protein structure based upon the Dictionary of Secondary Structure of Proteins (DSSP) assignment that defines eight structural states [46]. This has commonly been reduced to three central features; alpha-helices (H), extended beta-Strands (E), and coils (C) [47]. Assessment of the prediction accuracy of Hermes is carried out using Q3, the most widely used metric for measuring per-residue accuracy expressed as a percentage. An alternative to Q3, the Segment Overlap (SOV) score was developed to differentiate less critical boundary errors from those occurring inside structural segments and this was also employed [48].

## Results

### Optimising the architecture of Hermes

The structure of Hermes was controlled by three factors: (I) The number of well-established predictors in the protein layer; (II) the number of classifiers in the structure layer; (II) the complexity of the connection between the two. The simplest approach to a consensus predictor is a basic vote, whereby the most common prediction made by any number of predictors is taken as the final value. This was conducted for the top four predictors, Spider3, RaptorX-Property, SSpro and Porter4 and the top three, which removes Spider3. However, for the CASP12 dataset SSpro achieved anomalously high Q3/SOV results and so these were discarded. Hence, the simple consensus results used the next highest predictor and Hermes ran with only nine predictors. This low complexity solution was never more accurate than the predictors involved. Top four had Q3/SOV (%) accuracies for the blind test, CASP11, 12 and 13 of 89.0/84.6, 84.5/82.0, 79.6/72.3 and 85.4/79.5; top three of 93.3/90.6, 89.4/88.2, 82.9/76.2 and 86.9/86.2 (Table 1). Moreover, additional predictors to a simple vote reduces these Q3s despite adding additional information.

**Table 1.**
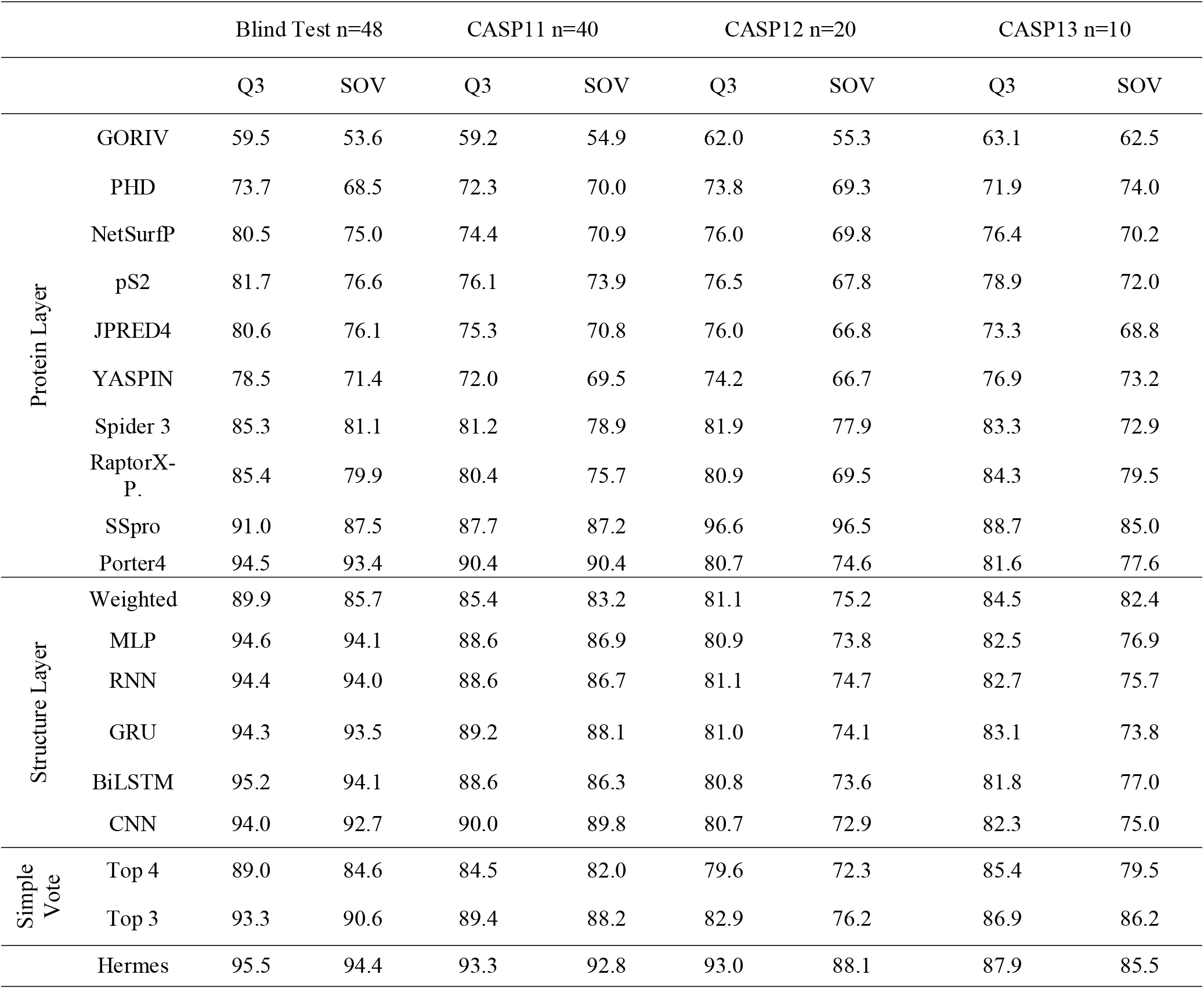
Accuracy and significance against Hermes for all Protein layer predictors

|  |  | Blind Test n=48 |  | CASP11 n=40 |  | CASP12 n=20 |  | CASP13 n=10 |  |
| --- | --- | --- | --- | --- | --- | --- | --- | --- | --- |
|  |  | Q3 | SOV | Q3 | SOV | Q3 | SOV | Q3 | SOV |
| Protein Layer | GORIV | 59.5 | 53.6 | 59.2 | 54.9 | 62.0 | 55.3 | 63.1 | 62.5 |
|  | PHD | 73.7 | 68.5 | 72.3 | 70.0 | 73.8 | 69.3 | 71.9 | 74.0 |
|  | NetSurfP | 80.5 | 75.0 | 74.4 | 70.9 | 76.0 | 69.8 | 76.4 | 70.2 |
|  | pS2 | 81.7 | 76.6 | 76.1 | 73.9 | 76.5 | 67.8 | 78.9 | 72.0 |
|  | JPRED4 | 80.6 | 76.1 | 75.3 | 70.8 | 76.0 | 66.8 | 73.3 | 68.8 |
|  | YASPIN | 78.5 | 71.4 | 72.0 | 69.5 | 74.2 | 66.7 | 76.9 | 73.2 |
|  | Spider 3 | 85.3 | 81.1 | 81.2 | 78.9 | 81.9 | 77.9 | 83.3 | 72.9 |
|  | RaptorX-P. | 85.4 | 79.9 | 80.4 | 75.7 | 80.9 | 69.5 | 84.3 | 79.5 |
|  | SSpro | 91.0 | 87.5 | 87.7 | 87.2 | 96.6 | 96.5 | 88.7 | 85.0 |
|  | Porter4 | 94.5 | 93.4 | 90.4 | 90.4 | 80.7 | 74.6 | 81.6 | 77.6 |
| Structure Layer | Weighted | 89.9 | 85.7 | 85.4 | 83.2 | 81.1 | 75.2 | 84.5 | 82.4 |
|  | MLP | 94.6 | 94.1 | 88.6 | 86.9 | 80.9 | 73.8 | 82.5 | 76.9 |
|  | RNN | 94.4 | 94.0 | 88.6 | 86.7 | 81.1 | 74.7 | 82.7 | 75.7 |
|  | GRU | 94.3 | 93.5 | 89.2 | 88.1 | 81.0 | 74.1 | 83.1 | 73.8 |
|  | BiLSTM | 95.2 | 94.1 | 88.6 | 86.3 | 80.8 | 73.6 | 81.8 | 77.0 |
|  | CNN | 94.0 | 92.7 | 90.0 | 89.8 | 80.7 | 72.9 | 82.3 | 75.0 |
| Simple Vote | Top 4 | 89.0 | 84.6 | 84.5 | 82.0 | 79.6 | 72.3 | 85.4 | 79.5 |
|  | Top 3 | 93.3 | 90.6 | 89.4 | 88.2 | 82.9 | 76.2 | 86.9 | 86.2 |
|  | Hermes | 95.5 | 94.4 | 93.3 | 92.8 | 93.0 | 88.1 | 87.9 | 85.5 |

To capture all available information, a diverse series of neural networks and a weighted vote were employed, which all act by different means to improve on a simple vote. However, none are able to consistently improve over the most accurate protein layer predicter; the best of these the BiLSTM achieved 95.2/94.1, 88.6/86.3, 80.8/73.6 and 81.8/77.0. This diverse information must be combined by the aggregator network, which achieved 95.5/94.4, 93.3/92.8, 93.0/88.1 and 87.9/85.5, greater than any input for all tests. We trained Hermes with every possible permutation of the ten Protein Layer predictors that contains SSpro and/or Porter4 (n=766), for example [GORIV, PHD, Spider3, Porter4] or [pS2, Spider3, SSpro, Porter4]. This was done using identical architecture (all eight structure layer predictors input to a final aggregator), regularisation, training and testing data. An increase in the number of Protein Layer predictors leads to a linear increase in Q3 accuracy on the blind test (Fig. 3a). Permutations were divided into three groups: (i) SSpro only [JPRED4, YASPIN, SSpro]; (ii) a Porter4 only [GORIV, PHD, Spider3, Porter4]; (iii) SSpro & Porter4 [Spider3, RaptorX-Property, SSpro, Porter4]. All permutations were superior to the most accurate input supplied, e.g. all permutations of [‘X’, SSpro] had a higher Q3 than that of SSpro. However, SSpro was consistently only ~3% less accurate than Porter4 alone, which was itself ~0.1% less accurate than SSpro & Porter4 (Fig. 3a). The most accurate architecture was all ten Protein layer predictors, including SSpro and Porter4. A maximum number of ten predictors were chosen for practical reasons to limit running times and dependencies on other servers.

**Fig. 3.**
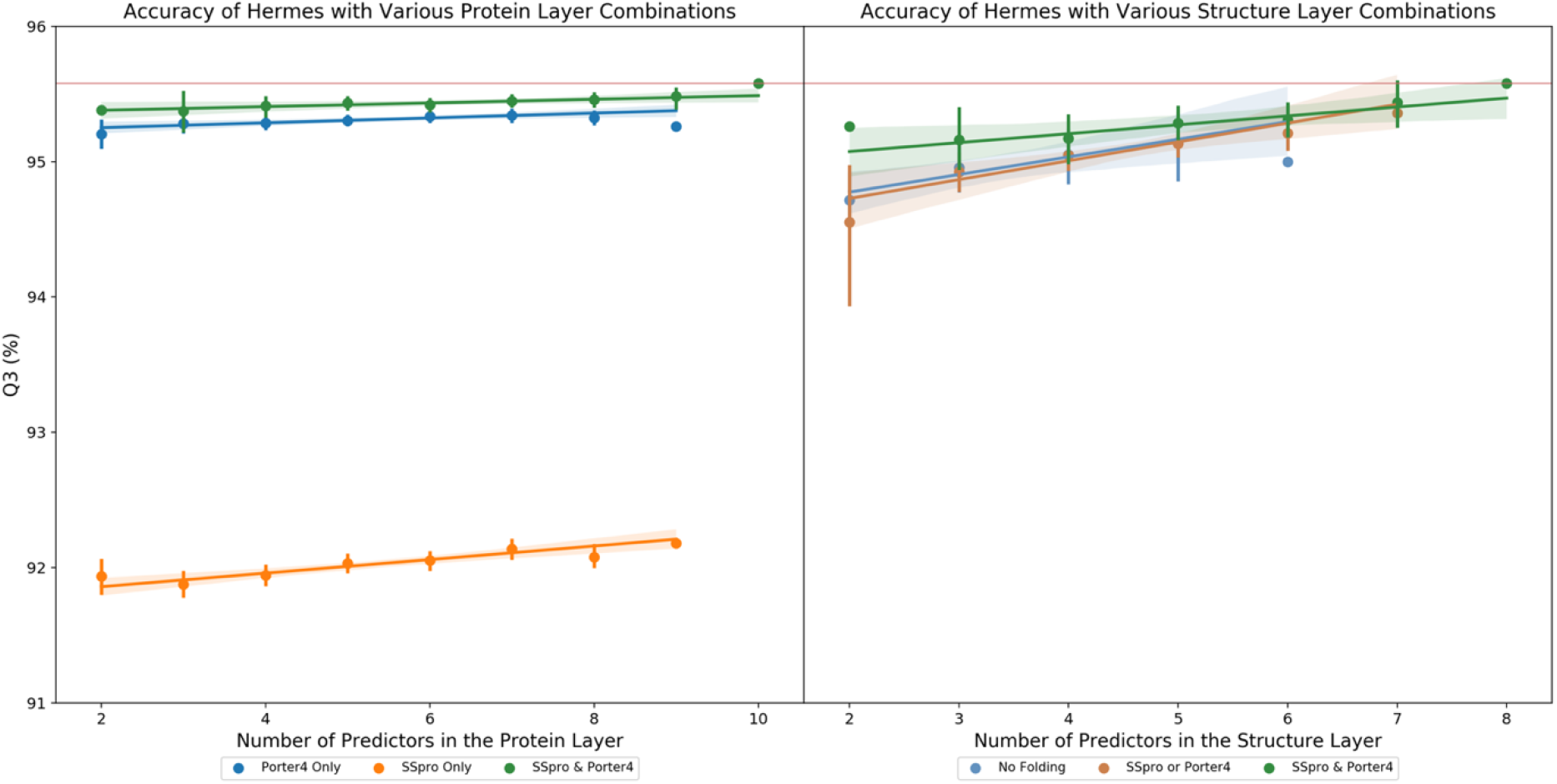
Q3 Accuracy by Hermes of Different Architecture. All training was performed on the JPRED training data and the identity reduced blind test: (a). The Q3 of predictions returned by Hermes with all possible permutations of the ten Protein layer predictors, that contain SSpro and/or Porter4, are shown. All other parameters used for the full Hermes pipeline were held constant. The data was divided into three groups containing; SSpro & Porter4, SSpro & not Porter4 and Porter4 & not SSpro. (b) The Q3 of predictions returned by Hermes with all possible permutations of the eight Structure layer predictors. Again, other variables were maintained and it was divided up into three groups; No Folding (Neither SSpro nor Porter4), SSpro or Porter4 and SSpro & Porter4. For both, (a&b). A horizontal red line indicates 95.5% prediction accuracy. This indicates the highest Q3 value, in both cases this was derived from the final Hermes architecture; ten Protein layer predictors and eight Structure layer predictors with folding of SSpro and Porter4.

Hermes was trained with all ten Protein layer predictors and every possible permutation of the eight Structure layer classifiers (n=246), for example [Weighted Vote, RNN, BiLSTM] or [GRU, CNN], again keeping all other variables constant. An increase in the number of Structure Layer predictors leads to a linear increase in blind test accuracy (Fig. 3b). We stratified these results into three groups: (i) no folding [Weighted Vote, MLP, RNN, GRU, BiLSTM, CNN]; (ii) a single fold with either SSpro or Porter4 [SSpro or Porter4, Weighted Vote, MLP, RNN, GRU, BiLSTM, CNN]; (iii) folding with both SSpro & Porter4 [SSpro, Porter4, Weighted Vote, MLP, RNN, GRU, BiLSTM, CNN]. The results of test ‘i’ showed a relatively steep gradient paralleling that of a single fold but with greater variance. Furthermore, in many permutations of test ‘ii’, structure layer classifiers were insufficient to improve on the most accurate Protein layer input. Test ‘iii’ was required to consistently surpass this mark.

Hermes with folding of either SSpro or Porter4 showed similar Q3 efficacy to no folding but with decreased variance (Fig. 3b). However, a sizeable increase in Q3 occurred when both SSpro & Porter4 were folded with the highest value occurring when all eight Structure layer predictors were used in conjunction. Moreover, this configuration led to the lowest variance. Notably, the variance displayed in Fig. 3a is generally greater than that of Fig. 3b. The stacking effect of ensemble learning is most efficacious with the complete array of Structure layer predictors. Thus, the architecture of Hermes shown here, featuring ten Protein layer predictors, eight Structure layer classifiers, and folding of SSpro and Porter4, was chosen for its balance between high Q3 performance on every dataset tested and its limiting of both dependencies and additional runtime.

### Hermes improves prediction accuracy and standard deviation

Hermes obtained a mean Q3 score of 95.5% ± 3.4 (Standard Deviation, STD) on the blind test. This is a considerable improvement over the accuracy of existing secondary structure predictors including: GORIV (59.5 ± 11.5), PHD (73.7 ± 9.6), NetSurfP (80.5 ± 10.2), pS2 (81.7 ± 10.3), JPRED4 (80.6 ± 7.7), YASPIN (78.5 ± 8.8), Spider3 (85.3 ± 7.1), RaptorX-Property (85.4 ± 7.5), SSpro (91.0 ± 8.4), and Porter4 (94.5 ± 4.3) (Table 1). Thus, Hermes oversaw, at minimum, a 1.0% improvement in Q3 over the current state-of-the-art template-based method (Porter4), and 10.1% over the most accurate *ab initio* method (RaptorX-Property). The smaller, hard-to-predict CASP11, 12 and 13 datasets caused reductions in accuracy for all ten Protein layer predictors. However, Hermes limited this Q3 degradation to 2.2, 2.5 & 7.6%. While Porter4 and RaptorX-Property decreased by 4.1, 13.8 & 12.9% and 5.0, 4.5 & 1.1%, respectively. Hermes shows diminishing returns, the lower the accuracy of the input data the greater the value Hermes can add. This is demonstrated by the average Q3 of the Protein Layer alongside the improvement made by Hermes, ordered as blind test, CASP13, 11 & 12: 81.0/14.5, 77.8/10.06, 76.9/16.4, 75.8/17.2%.

In the 48 blind test predictions, the mean updated Q3 of each successive rolling prediction remained higher with Hermes than with other widely used models and reached stability earlier (Fig. 4b). For the end user, a single mean Q3 is uninformative if the query sequence will likely fail to attain it. In this case, variance is a more useful statistic. Hermes shows significantly reduced standard deviation, compared to all Protein layer predictors (Fig. 4c). In the quartile Q3 distribution of blind test proteins the minimum Q3 of Hermes, 88.2%, is greater than the respective means of all ten Protein layer predictors, with the exception of SSpro, and Porter4 (Fig. 4c). Moreover, these two classifiers each have 2 lower outliers compared to the 1 of Hermes. Histograms of the blind test Q3 spread for Hermes, Porter4, and all Protein layer predictors are all normally distributed, as measured by Shapiro-Wilk test with p-values = 7.3×10^−4^, 2.0×10^−3^, 1.1×10^−12^, respectively (Fig. 4a). Direct comparison with all models revealed that Hermes is greater than or equal to all Protein layer predictors (pairwise) at ≥95.4% of blind test proteins, excluding SSpro and Porter4, which are 83.3% and 79.2%, respectively (Table 2).

**Table 2.** The Percentage of outcomes in which Hermes achieved the greatest Q3 score

|  | < Hermes (%) | = Hermes (%) | > Hermes (%) | ≤ Hermes (%) |
| --- | --- | --- | --- | --- |
| GORIV | 100.0 | 0.0 | 0.0 | 100.0 |
| PHD | 100.0 | 0.0 | 0.0 | 100.0 |
| NetSurfP | 100.0 | 0.0 | 0.0 | 100.0 |
| pS2 | 95.8 | 4.2 | 0.0 | 100.0 |
| JPRED4 | 97.9 | 2.1 | 0.0 | 100.0 |
| YASPIN | 97.9 | 2.1 | 0.0 | 100.0 |
| Spider 3 | 91.7 | 0.0 | 8.3 | 91.7 |
| RaptorX-Property | 93.8 | 2.1 | 4.2 | 95.8 |
| SSpro | 70.8 | 12.5 | 16.7 | 83.3 |
| Porter4 | 45.8 | 33.3 | 20.8 | 79.2 |

**Fig. 4.**
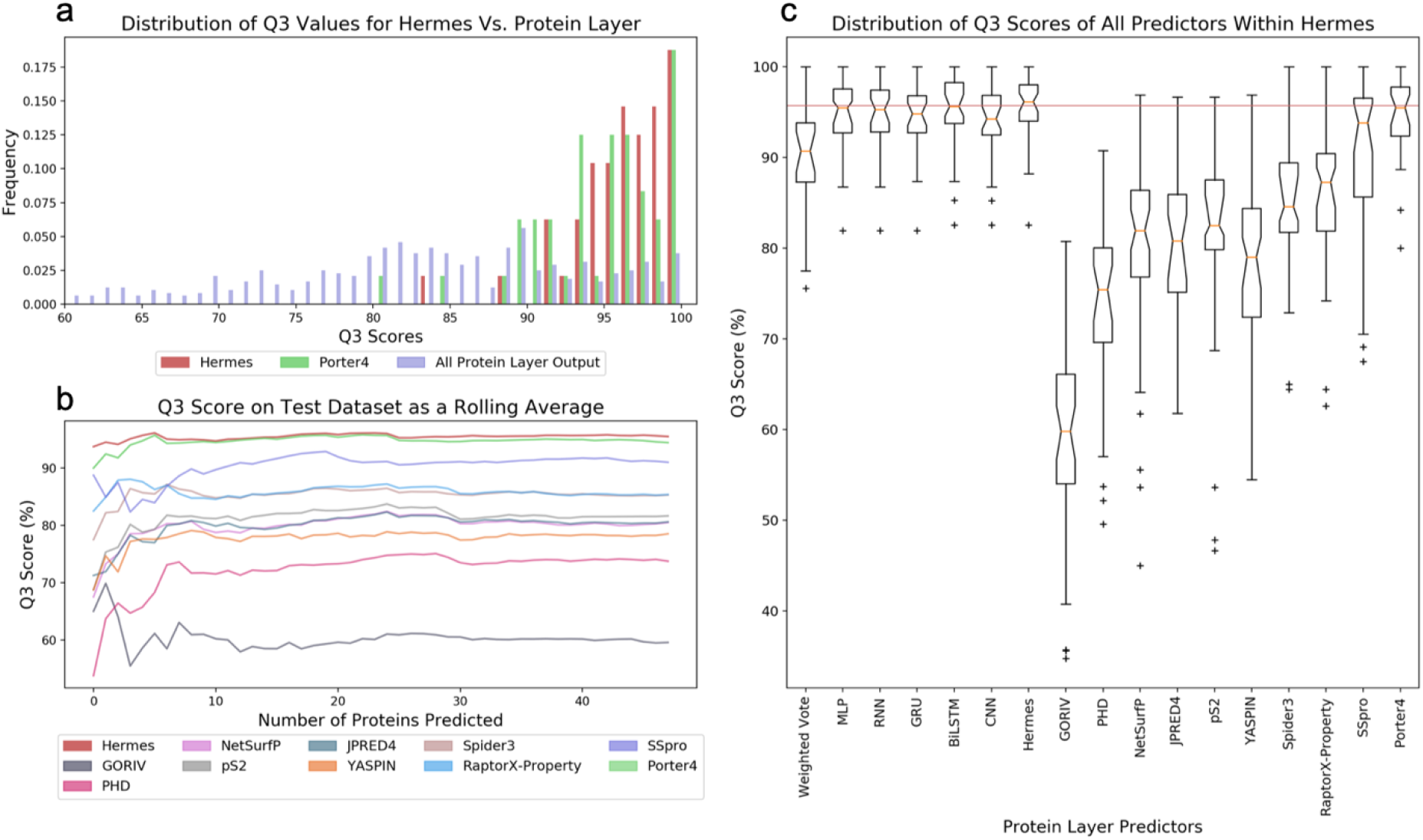
Analysis of error and value added by Hermes as a rolling average and distribution of Q3 Scores. All data was collected and calculated from the JPRED blind test, the following plots are shown: (a) The normalised frequency of Q3 scores is displayed for Hermes, Porter4, and all of the ten Protein layer classifiers, including Porter4. For the former two predictors ten copies of the collected data is plotted, thus all variables have the same quantity of data and frequencies are normalised accordingly. (b) A rolling average of Q3 scores was calculated as the updated mean Q3 for the run with every additional protein predicted, for all ten Protein layer predictors. (c) The distribution of Q3 scores are shown as a boxplot for all sixteen predictors within Hermes and the final output.

### Hermes is accurate at predicting all three DSSP states

The 7-Fold Cross Validation (CV) trained models returned prediction accuracies of 97.3, 97.5, 97.0, 97.7, 97.1, 97.0, and 95.5%, with a mean 97.0% ± 0.66 (STD). The Q3 of Hermes decreases by 1% with every additional 53 residues of query protein length (Fig. 5a). Hence, for a modal PDB protein of about 350 residues an accuracy of 91.1 could be expected. A confusion matrix of the predicted states (H, E, and C) plotted against experimentally derived states for the blind test (Fig. 5b) revealed the proportion of predictions that were either correct (Q3_**H**_, Q3_**E**_, Q3_**C**_ with Q3 of 0.93, 0.98, and 0.95 respectively) or incorrect. While, all were within 2.5% of the Hermes mean Q3, coiled residues (Q3_**C**_) were the lowest performing state, with 3.0% being mislabelled as an alpha-helix. The most accurate predictions were for extended beta-strands, a DSSP class that has traditionally been the most difficult to predict correctly [49–51]. Furthermore, a detailed view of beta-strands showed that these mistakes occurred especially where the residue is Proline, Cysteine, and Glycine, giving accuracies of 94.1, 94.9, and 95.3%, respectively (Fig. 5c). Proline and Cysteine can be explained by their actions as alpha-helix breakers and the formation of disulphide bonds, respectively. Notably, Aspartate has an accuracy of 96.6%, while separated by just a single methyl group, Glutamate, has a very similar Q3 of 96.2%, suggesting that in this analysis there may be direct biological *in-silico* correlations.

**Fig. 5.**
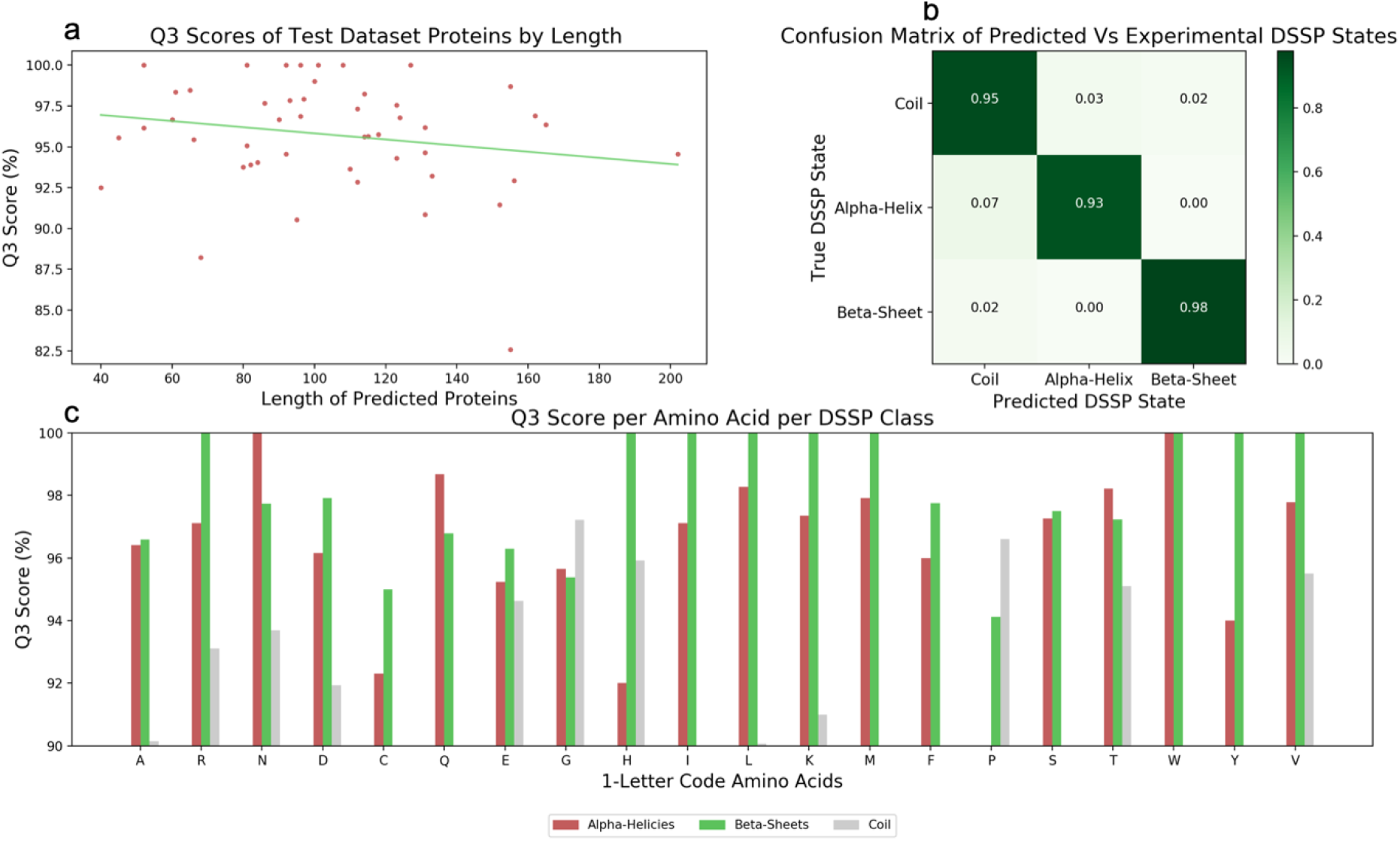
An Analysis of the errors within Hermes. Data was collected and calculated from the JPRED blind test for predictions made solely by Hermes. The following are illustrated: (a) A scatter plot for the Q3 scores against the length of the predicted protein from which it was derived. A linear regression is displayed in green with equation y = −0.019(x) + 97.70; (b) A confusion matrix showing the percentages of correctly predicted secondary structure states, given by DSSP. (c) The Q3 scores from the 48 secondary structure predictions of Hermes, for each of the twenty standard amino acids and then for each of the three DSSP states (H, E, C).

### Hermes prediction accuracy is robust to sparse input data

The homology and sparsity of available templates is not correlated with Hermes’ prediction accuracy. Data sparsity, where a PSI-BLAST search for templates yields a limited number of poor match’s, can impact a predictors accuracy as there are no high quality solutions [52]. We compared the impact of sparsity via PSI-BLAST searches (3 iterations at E value = 0.001) of the non-redundant database for both the JPRED blind test dataset and the CASP11 dataset (Fig. 6 a-b). A directly proportional relationship between Q3 and mean identity, calculated as the average identity of all retrieved hits for each proteins’ MSA, would indicate that the identity of putative templates plays a significant role in prediction accuracy (Fig. 6a). High Q3 values were obtained across the range of mean identities for both datasets, despite CASP11 accuracies being generally lower (Fig. 6a). Pearson’s correlation coefficients of −0.13 and −0.079 were obtained for the blind test and CASP11. For example, 98% and 100% Q3 scores for both occurred at identities of 21 and 26%, respectively. Such disparity in numbers indicates that accuracy, for these predictions, is not afforded by templates. Mean identity of an MSA is a more relevant measure than maximum identity because proteins are often fragmented in template-based methods.

**Fig. 6.**
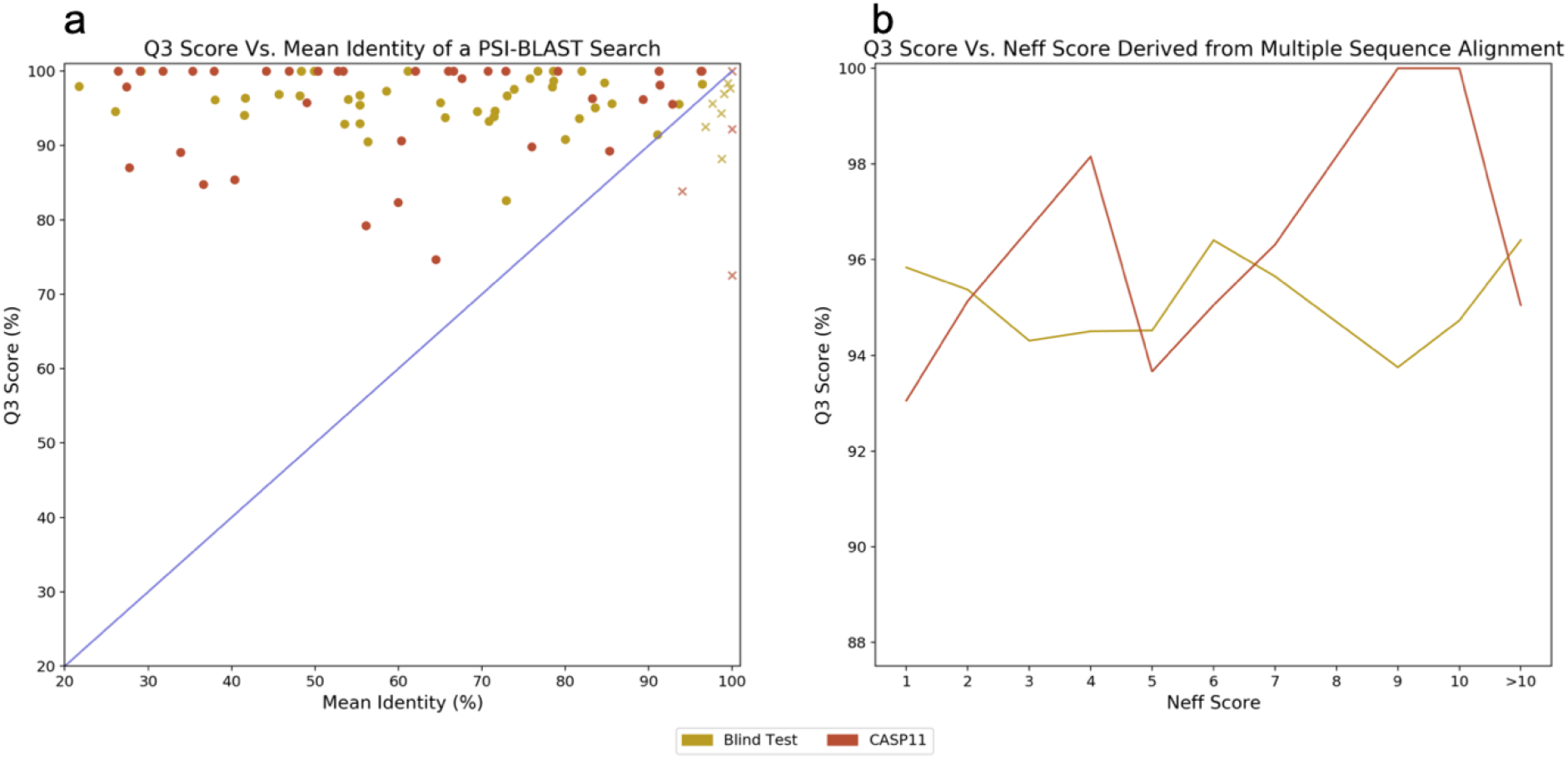
Sparsity of protein layer inputs. Hermes predictions were made on the identity reduced CASP11 dataset (yellow), and the blind test dataset (red). (a) For the proteins of both datasets, the mean identity of PSI-BLAST MSAs are plotted with the Q3 scores from predictions of those proteins. Homology searches were conducted with E value = 0.001, on the non-redundant database, for 3 iterations. Shown as an X Y scatter, with a blue y=x line to represent direct proportionality. Q3 scores greater than the respective identity values have circular markers, while Q3 accuracies below the respective identity values have cross markers. (b) Q3 scores are plotted against the Number of Effective (NEFF) sequences scores, a value from 1-20, although both datasets had a maximum NEFF=16. The NEFF at a column of the MSA is calculated via exp(−∑_a_ p_a_ ln p_a_), this value is then averaged across all columns. Values are shown as integer bins, with the means plotted as lines.

Accuracy can also be compared to the number of effective sequence homologs (NEFF) to determine the impact of data sparsity [53,54]. The NEFF of the protein can be considered as the average Shannon entropy of a MSA, and is given by exp(−∑*_a_ p_a_* ln *p*_*a*_) [55]. The result is a real number, indicating a point for each unique amino acid of the 20 naturally occurring (Fig. 6b), with a probability of finding each unique amino acid at a given NEFF score. A higher score indicates a richer MSA, that likely possess greater homology information. In analysis such as this, a convex curve would illustrate how limited Q3 accuracy corresponds with low NEFF, often increasing to a plateau (around NEFF=10). Typically, the accuracy gap between NEFF=1 and the plateau is 8-10% [25]. However, on both datasets, Hermes maintained accuracy across the NEFF range. The maximum Q3 spread between any two points, on the blind test, reduced to just 2.6%, with the lowest value at the NEFF=9. CASP11 predictions maintained independency from NEFF, albeit with a greater maximum Q3 spread at 7.0%. The functionality of Hermes therefore preserves accuracy without complete reliance on homology.

The combination of Protein layer predictors controls the distribution of DSSP states available for optimisation by Hermes. Predictor selection was largely based on two key criteria, congruence and accuracy. It is possible to outline these metrics for the final set of Protein layer predictors by comparing the congruence, or agreement, of each predictor to every other predictor on the blind test (Fig. 7a). Two predictors of similar accuracy may share relatively little congruence, and thus be suited for ensemble meta-learning. Whereas the reverse, high congruence, should be avoided because many copies of the same data acts to increase noise. Template-based Porter4 and SSpro methods are the most similar pair, sharing a proportion of 0.94 predictions, undoubtedly because they use many of the same templates. The recently updated Spider3 and RaptorX-Property protocols share 0.89. NetSurfP, JPRED4, and pS2 form a triplet of *ab-initio* and older template-based methods with a congruence of 0.86-0.88. Finally, there are three singleton groups GORIV, PHD, and YASPIN. The lack of congruence for GORIV and PHD with every other predictor is unsurprising as they represent a roughly 20-year disparity in research. Thus, the singletons supply informed variability to the information passed, without incorporating excessive inaccurate noise. Frequency of occurrence, i.e. how frequently a certain number of DSSP states arises in each 10-element array, was found to be bimodal (Fig. 7c). The majority of arrays feature 100% congruence, whereby all 10 elements are the same DSSP state. Further, the frequency of arrays then declines from 10 congruent elements to 9 to 8 and plateaus at 7 to 5. This trend is maintained for all three DSSP states, suggesting similar aptitude at predicting each. Thus, we may infer that congruence is highly clustered, that is to say, predictors agree completely for the majority of arrays but disagree extensively in a minority (Fig. 7). This later case likely being ambiguous or hard-to-predict amino acids that could reasonably be assigned differing states. For each predictors, the number of predicted states was compared to the number of experimental states (Fig. 7b). Except for SSpro, no predictor was accurate on all three DSSP states. For example, Porter4 was accurate at alpha-helices, but over-predicted coiled states and under-predicted Beta sheets. Each predictor had advantages for stretches of X states, when flanked by Y and Z states. The significant improvements of Hermes manifest by learning these pattern-specific advantages and disadvantages during training to apply an appropriate weight for each pattern, per predictor.

**Fig. 7.**
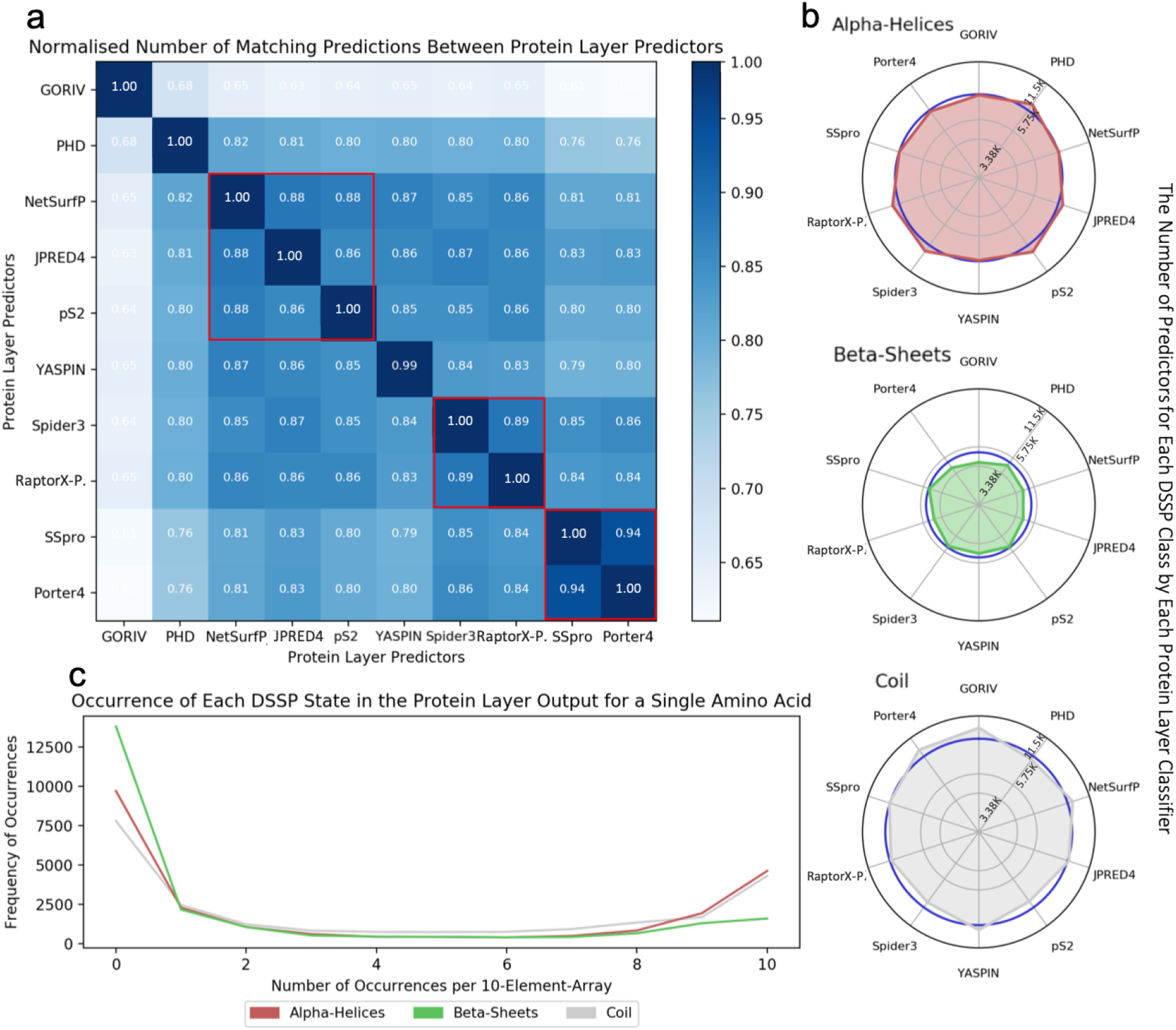
Sparsity of structure layer inputs. Data was collected and calculated from the JPRED blind test for predictions made by the ten Protein layer predictors, the following are shown: (a) A 10×10 density matrix of congruence, as measured by matching DSSP predictions, between each predictor in the Protein layer to every other predictor in the same layer. Red boxes indicate predictors with congruence ≥ 0.89. (b) Three spider diagrams, one for each DSSP state, shows the number of predicted states made by each predictor. Blue circles represent the true, experimentally derived, number of DSSP states. (c) A line graph of congruence within every 10-element array present in the dataset. The frequency of occurrence, for each level of congruence, is plotted against the number of matching predictions per array, per state.

## Discussion

The principal advantages of Hermes come from amalgamating the most successful previous research efforts to elucidate relationships, not in amino acids, but within meta structural information. Hermes has a mean Q3 accuracy of 95.5% and SOV of 94.4%, following the rigorous JPRED4 evaluation regime. We have made available the runnable source code, predictions and their accuracies, and the cross-validation models for Hermes. It has long been known that a validated Q3 of 100% is unattainable due to theoretical limits, as secondary structure is purely nominal. Tertiary and quaternary structure, and environmental interactions, all affect local conformations, that is to say, until we can see the entire protein we will be relegated to predicting from constituent parts. Hermes can run locally in minutes on a standard computer, while presenting minimal variance, and limiting accuracy depreciation on hard-to-predict targets. This has potential to provide confidence to the end-user that their queries can attain quoted accuracies, at low cost, and be reliably used to draw single-base-resolution conclusions.

Hermes is a versatile architecture that can optimise for problems previously attempted, but not yet solved, by exploring and learning meta-information. The integration of various approaches as an ensemble network, with care taken to exploit advantages and limit disadvantages, is a robust way to advance the current state of prediction research. As such, more than sixty years of high quality research in the field has granted a repertoire of different implementations that, when combined, offer enhanced solutions. Diversion of the Hermes pipeline for protein contact prediction, or by using another comparable predictor, coupled with the secondary structure work presented here, can be fed into the CONFOLD pipeline for 3D tertiary structure models [56]. While *in silico* 3D protein structure prediction remains a relatively new technique, integration with the Hermes pipeline provides inputted data of the greatest fidelity. As protein structure prediction become more accurate it may be possible to optimise pipelines such as Hermes for optimization of 8-state secondary structure prediction, and consider the development of methodology for tertiary prediction.

## References

1. Terwilliger TC, Stuart D, Yokoyama S. Lessons from Structural Genomics. Annual Review of Biophysics. 2009;38:371–83.

2. Fasman GD, Chou PY. Conformational parameters of amino acids in helical, beta-sheet and random coil regions calculated from proteins. Biochemistry. 1974;13:211–22.

3. Garnier J, Osguthorpe DJ, Robson B. Analysis of the accuracy and implications of simple methods for predicting the secondary structure of globular proteins. Journal of Molecular Biology. 1978;120:97–120.

4. Garnier, J. and Robson B. Prediction of Protein Structure and the Principles of Protein Conformation. Prediction of Protein Structure and the Principles of Protein Conformation. 1989.

5. Gibrat JF, Garnier J, Robson B. Further Developments of Protein Secondary Structure Prediction Using Information Theory New Parameters and Consideration of Residue Pairs. J mol Biol. 1987;198:425–43.

6. Garnier J, Gibrat JF, Robson B. GOR secondary structure prediction method version IV. Methods Enzymol. 1996;266:540–53.

7. Sen TZ, Jernigan RL, Garnier J, Kloczkowski A. GOR V server for protein secondary structure prediction. Bioinformatics. 2005;21:2787–8.

8. Cheng J. A multi-template combination algorithm for protein comparative modeling. BMC Structural Biology. 2008;8.

9. Bondugula R, Xu D. MUPRED: A tool for bridging the gap between template based methods and sequence profile based methods for protein secondary structure prediction. Proteins: Structure, Function and Bioinformatics. 2006;66:664–70.

10. Xu D, Bondugula R. MUPRED: A tool for bridging the gap between template based methods and sequence profile based methods for protein secondary structure prediction. Proteins: Structure, Function and Bioinformatics. 2007;66:664–70.

11. Pollastri G, Martin AJ, Mooney C, Vullo A. Accurate prediction of protein secondary structure and solvent accessibility by consensus combiners of sequence and structure information. BMC Bioinformatics. 2007;8:201.

12. Zickmann F, Renard BY. IPred - integrating ab initio and evidence based gene predictions to improve prediction accuracy. BMC Genomics. 2015;16.

13. Mistry J, Kloppmann E, Rost B, Punta M. An estimated 5% of new protein structures solved today represent a new Pfam family. Acta Crystallographica Section D: Biological Crystallography. 2013. p. 2186–93.

14. Zhang W, Dunker AK, Zhou Y. Assessing secondary structure assignment of protein structures by using pairwise sequence-alignment benchmarks. Proteins: Structure, Function and Genetics. 2008;71:61–7.

15. Rost B. Twilight zone of protein sequence alignments. Protein Engineering. 1999;12:85–94.

16. Sagi O, Rokach L. Ensemble learning: A survey. Wiley Interdisciplinary Reviews: Data Mining and Knowledge Discovery. 2018;e1249.

17. Rokach L. Ensemble-based classifiers. Artificial Intelligence Review. 2010;33:1–39.

18. Bell R, Koren Y, Volinsky C. The BellKor 2008 Solution to the Netflix Prize. Netflix prize documentation. 2009;1–21.

19. Wolpert DH. Stacked generalization. Neural Networks. 1992;

20. Rost B. PHD: predicting 1D protein structure byprofile based neural networks. Methods in Enzymol. 1996;266:525–39.

21. Petersen B, Petersen TN, Andersen P, Nielsen M, Lundegaard C. A generic method for assignment of reliability scores applied to solvent accessibility predictions. BMC Structural Biology. 2009;9.

22. Drozdetskiy A, Cole C, Procter J, Barton G. JPred 4□: a protein secondary structure prediction server. Nucleic Acids Res. 2015;43:389–94.

23. Lin K, Simossis V a, Taylor WR, Heringa J. A simple and fast secondary structure prediction method using hidden neural networks. Bioinformatics (Oxford, England). 2005;21:152–9.

24. Heffernan R, Yang Y, Paliwal K, Zhou Y. Capturing non-local interactions by long short-term memory bidirectional recurrent neural networks for improving prediction of protein secondary structure, backbone angles, contact numbers and solvent accessibility. Bioinformatics. 2017;33:2842–9.

25. Wang S, Peng J, Ma J, Xu J. Protein Secondary Structure Prediction Using Deep Convolutional Neural Fields. Scientific Reports. 2016;6.

26. Chen CC, Hwang JK, Yang JM. (PS)2-v2: Template-based protein structure prediction server. BMC Bioinformatics. 2009;10:366.

27. Magnan CN, Baldi P. SSpro/ACCpro 5: Almost perfect prediction of protein secondary structure and relative solvent accessibility using profiles, machine learning and structural similarity. Bioinformatics. 2014;30:2592–7.

28. Mooney C, Pollastri G. Beyond the Twilight Zone: Automated prediction of structural properties of proteins by recursive neural networks and remote homology information. Proteins: Structure, Function and Bioinformatics. 2009;77:181–90.

29. White BW, Rosenblatt F. Principles of Neurodynamics: Perceptrons and the Theory of Brain Mechanisms. The American Journal of Psychology. 1963;76:705.

30. Miljanovic M. Comparative analysis of Recurrent and Finite Impulse Response Neural Networks in Time Series Prediction. Indian Journal of Computer Science and Engineering (IJCSE). 2013;

31. van Merrienboer B, Bahdanau D, Bougares F, Schwenk H, Bengio Y. Learning Phrase Representations using RNN Encoder-Decoder for Statistical Machine Translation. Proceedings of the 2014 Conference on Empirical Methods in Natural Language Processing (EMNLP). 2014. p. 1724–34.

32. Hochreiter S, Urgen Schmidhuber J. LONG SHORT-TERM MEMORY. Neural Computation. 1997;9:1735–80.

33. LeCun Y, Bottou L, Bengio Y, Haffner P. Gradient-based learning applied to document recognition. Proceedings of the IEEE. 1998;86:2278–323.

34. Parvin H, Mirnabibaboli M, Alinejad-Rokny H. Proposing a classifier ensemble framework based on classifier selection and decision tree. Engineering Applications of Artificial Intelligence. 2015;37:34–42.

35. Barton G, Drozdetskiy A. JPred 4: JNet training (v.2.3.1) details.

36. Moult J, Fidelis K, Kryshtafovych A, Schwede T, Tramontano A. Critical assessment of methods of protein structure prediction: Progress and new directions in round XI. Proteins: Structure, Function and Bioinformatics. 2016;

37. Moult J, Fidelis K, Kryshtafovych A, Schwede T, Tramontano A. Critical assessment of methods of protein structure prediction (CASP)—Round XII. Proteins: Structure, Function and Bioinformatics. 2018;

38. Heffernan R, Paliwal K, Lyons J, Dehzangi A, Sharma A, Wang J, et al. Improving prediction of secondary structure, local backbone angles, and solvent accessible surface area of proteins by iterative deep learning. Scientific Reports. 2015;5.

39. Kline DM, Berardi VL. Revisiting squared-error and cross-entropy functions for training neural network classifiers. Neural Computing and Applications. 2005;14:310–8.

40. Pourtaheri ZK, Zahiri SH. Ensemble classifiers with improved overfitting. 1st Conference on Swarm Intelligence and Evolutionary Computation, CSIEC 2016 - Proceedings. 2016. p. 93–7.

41. Prechelt L. Early stopping - But when? Lecture Notes in Computer Science (including subseries Lecture Notes in Artificial Intelligence and Lecture Notes in Bioinformatics). 2012;7700 LECTU:53–67.

42. Srivastava N, Hinton G, Krizhevsky A, Sutskever I, Salakhutdinov R. Dropout: A Simple Way to Prevent Neural Networks from Overfitting. Journal of Machine Learning Research. 2014;15:1929–58.

43. Ng AY. Feature selection, L1 vs. L2 regularization, and rotational invariance. Twenty-first international conference on Machine learning - ICML’04. 2004;78.

44. Bergstra J, Bardenet R, Bengio Y, Kégl B. Algorithms for Hyper-Parameter Optimization. Advances in Neural Information Processing Systems (NIPS). 2011. p. 2546–54.

45. Abadi M, Barham P, Chen J, Chen Z, Davis A, Dean J, et al. TensorFlow: A System for Large-Scale Machine Learning TensorFlow: A system for large-scale machine learning. 12th USENIX Symposium on Operating Systems Design and Implementation (OSDI’16). 2016. p. 265–84.

46. Kabsch W, Sander C. Dictionary of protein secondary structure: Pattern recognition of hydrogen□bonded and geometrical features. Biopolymers. 1983;22:2577–637.

47. Kim H, Park H. Protein secondary structure prediction based on an improved support vector machines approach. Protein engineering. 2003;16:553–60.

48. Zemla A, Venclovas Č, Fidelis K, Rost B. A modified definition of Sov, a segment-based measure for protein secondary structure prediction assessment. Proteins: Structure, Function and Genetics. 1999;34:220–3.

49. Zhang H, Zhang T, Chen K, Kedarisetti KD, Mizianty MJ, Bao Q, et al. Critical assessment of high-throughput standalone methods for secondary structure prediction. Briefings in Bioinformatics. 2011;12:672–88.

50. Crivelli S, Eskow E, Bader B, Lamberti V, Byrd R, Schnabel R, et al. A physical approach to protein structure prediction. Biophysical Journal. 2002;82:36–49.

51. Kloczkowski A, Ting KL, Jernigan RL, Garnier J. Combining the GOR V algorithm with evolutionary information for protein secondary structure prediction from amino acid sequence. Proteins: Structure, Function and Genetics. 2002;49:154–66.

52. Yang JY, Peng ZL, Chen X. Prediction of protein structural classes for low-homology sequences based on predicted secondary structure. BMC Bioinformatics. 2010;11.

53. Ma J, Peng J, Wang S, Xu J. A conditional neural fields model for protein threading. Bioinformatics. 2012;28.

54. Orlando G, Raimondi D, Vranken WF. Observation selection bias in contact prediction and its implications for structural bioinformatics. Scientific Reports. 2016;6.

55. Xu J, Wang S, Ma J. Protein Homology Detection Through Alignment of Markov Random Fields: Using MRFalign. 2015.

56. Adhikari B, Bhattacharya D, Cao R, Cheng J. CONFOLD: Residue-residue contact-guided ab initio protein folding. Proteins: Structure, Function and Bioinformatics. 2015;83:1436–49.

